# Age-Dependent Mechanisms of Cardiac Hypertrophy Regression Following Exercise in Female Mice

**DOI:** 10.1101/2025.04.07.647563

**Authors:** Leslie A. Leinwand, Claudia Crocini

## Abstract

Cardiac adaptation to exercise is a fundamental physiological process, but its regression and the underlying molecular mechanisms, particularly in relation to age, remain poorly understood. This study investigated the age-dependent differences in cardiac remodeling and molecular signaling during exercise training and detraining in young (5-week-old) and adult (24-week-old) female mice, focusing specifically on how cardiac plasticity changes with adulthood rather than senescence. While both age groups exhibited significant cardiac hypertrophy after the exercise period, young mice displayed significantly more hypertrophic growth (23% increase in left ventricular mass versus 15% in adults). During detraining, cardiac mass regression occurred more rapidly in young mice. Transcriptomic analysis revealed distinct gene expression profiles between age groups, with changes in metabolic and autophagy pathways. Notably, ERK1/2 phosphorylation increased significantly during exercise in young but not adult hearts, correlating with elevated expression of well-known genes associated with exercise, namely CITED4 and SOD2. Furthermore, increased LC3-II/LC3-I ratio and AMPK phosphorylation were observed exclusively in young mice during detraining, indicating age-specific activation of autophagy-mediated cardiac remodeling. These findings demonstrate that cardiac adaptability to exercise and detraining follows distinct molecular pathways in young versus adult mice, with the younger heart exhibiting greater plasticity through enhanced ERK signaling during hypertrophy and autophagy during regression. This age-dependent cardiac plasticity may have important implications for understanding the cardiovascular benefits of exercise across the lifespan and developing age-appropriate exercise recommendations.

## INTRODUCTION

The heart demonstrates remarkable plasticity, adapting its size and function to several stimuli, including exercise^1^. Exercise-induced cardiac remodeling is characterized by increased cardiac mass, which can quickly regress during periods of reduced physical activity or detraining^2^. This process of cardiac adaptation and regression has been observed in both humans and animal models^1,3,4^. Rodents, especially, have been extensively employed to study cardiac hypertrophy in response to exercise. Three types of rodent exercise training protocols dominate the literature: treadmill running, swim training, and voluntary cage wheel running. Although a conditioning phase is usually included in swim and treadmill exercise protocols, the nature of these types of exercise remains forced and may involve physiologic stress in the animals^5^. Conversely, voluntary cage wheel running mimics the natural tendencies of rodents to move and it is performed in total autonomy during the regular wake hours for the animals. Regardless of the type of exercise, numerous studies (reviewed in ^6^) have documented molecular mechanisms associated with cardiac growth, e.g. activation of the phosphoinositide 3-kinase (PI3K)/Akt^7^ and the extracellular signal-regulated kinase (ERK1/2) pathways^8^, increased levels of the CBP/p300-interacting transactivator with ED-rich carboxy-terminal domain-4 (CITED4) downstream of ERK^9^, metabolic reprogramming with increased antioxidant defenses, including superoxide dismutase 2 (SOD2)^10^. In contrast, pathological cardiac hypertrophy induced, for example, by aortic constriction results in the increased expression of atrial or brain natriuretic peptide (ANP, BNP) and β-myosin heavy chain^7^.

Additionally, genetic background, sex, and age are all important modifiers of exercise performance and exercise-induced cardiac hypertrophy in rodents. An extensive study employing 7 mouse strains showed significant differences in average running distance and speed among the different strains with voluntary wheel running^11^. This study used male mice for all strains and demonstrated some differences in running performance and cardiac adaptation to exercise. Other studies have indicated that generally male mice show a mild cardiac response to exercise, with 5.0 ± 1.6% cardiac mass increase in C57B6/J and 9.0 ± 2.2% in FVB/NJ background^12^. However, a significant sex dimorphism has been demonstrated in exercise-induced cardiac remodeling in animal models. Female mice and rats demonstrate more pronounced hypertrophic responses compared to their male counterparts^12–15^. Depending on the strain, female mice showed about 15 to 25% cardiac mass increase with 21-days of voluntary wheel running. Age-related differences in cardiac plasticity have also been observed in both preclinical and clinical settings. While training in young animals generally induced some degree of cardiac hypertrophy, exercised-induced cardiac growth in aged animals is remarkably variable^16–21^ and it can be even paradoxically reduced^22^. In humans, younger individuals often display greater adaptive responses to exercise, whereas older adults exhibit attenuated cardiac reserve^23,24^, i.e. ability to increase cardiac output to meet the higher demands of exercise. Studies suggested that this difference may stem from age-dependent variations in signaling pathways governing cardiac hypertrophy, from Ca^2+^ handling, to signaling pathways and metabolism (reviewed in^23^). Obviously, most studies have focused on the beneficial role of exercise in attenuating aging in healthy and diseased patients. Studies comparing sedentary and athletic older adults have suggested that lifelong physical activity is associated with less cardiac aging^25^. Exercise is also proposed as therapeutic measure in certain cardiac diseases, such as heart failure with preserved ejection fraction (HFpEF) and other cardiometabolic diseases^26^.

The fact that exercise-induced hypertrophy is physiological and can fully regress is a fundamental principle. When the exercise stimulus ceases, the heart typically returns to its pre-training dimensions rapidly. For example, cardiac hypertrophy induced by 21 days of swimming in C57B6/J male mice regressed within 1 week^27^. However, the molecular mechanisms responsible for regression of cardiac hypertrophy upon cessation of exercise activity have been much less studied (reviewed in^28^. It is expected that the reduction of the exercise-induced hypertrophic signaling *per se* could underlie the cardiac mass regression. However, considering the fast time-frame of regression, active remodeling cannot be excluded. Autophagy, a cellular process for degrading and recycling cellular components, may be directly involved to degrade cardiac muscle proteins. Autophagy can be activated by several pathways, including the AMP-activated protein kinase (AMPK) pathway, a key regulator of cellular energy homeostasis. The role of age in regression from exercise-induced cardiac hypertrophy has not been investigated.

The present work aimed to study the age-dependent differences in cardiac remodeling during exercise training and detraining in adulthood using young (5-week-old) and adult (24-week-old) mice. By combining transcriptomic and protein analyses, we sought to elucidate the molecular mechanisms underlying the differential cardiac responses to exercise and detraining across age groups. Understanding these age-specific adaptations may provide insights into optimizing exercise recommendations and timing across the lifespan.

## METHODS

### Animals and exercise program

All animal treatments were approved by the Institutional Animal Care and Use Committee at the University of Colorado Boulder (Protocol #2351) and are in accord with the NIH guidelines. Wild-type, 5 or 24 week-old C57Bl/6 male and female mice (Jackson Laboratories) were fed *ad libitum* standard rodent chow and housed in a 12-hour light/dark cycle.

### Total protein extraction

Frozen left ventricular tissue samples were homogenized and solubilized in Urea-Thiourea sample buffer (8M Urea, 2M Thiourea, 50mM Tris (pH 6.8), 75mM DTT, 3% SDS, 0.05% Bromophenol Blue)^29^ at the ratio of 1:10. Samples were vortexed, centrifuged at 14,500 rpm for 8 minutes at 4° C. The supernatant was collected and aliquots frozen and stored at -80°C. Protein concentration was determined using Pierce 660nm reagent (ThermoFisher). Prior to gel loading, samples were thawed, heated for 4 minutes at 55°C, vortexed and centrifuged at maximum speed for a minute.

### Western blots

Equal concentrations (0.5 µg/µl) of protein were resolved by SDS-Page using 4-12% NuPAGE™ Novex™ 4-12% Bis-Tris Protein Gels and MES SDS Running Buffer (Thermo Fisher). Proteins were transferred to nitrocellulose membranes (Millipore Corp., Billerica, MA, USA) for immunoblot analysis. For immunoblot analysis, membranes were blocked with 5% bovine serum albumin (BSA) in Tris-buffered saline buffer containing 0.1% of Tween (Fisher) (TBS-T). Membrane were incubated with primary antibodies in 5% BSA in TBS-T at 4°C, on a shaker, overnight. Membranes were then washed and incubated-with secondary antibodies in 2.5% milk powder in TBS-T. Secondary antibodies were: HRP-goat anti rabbit (Cell Signaling Technology #7074) and HRP-goat anti mouse (Jackson # 115-035-003). Blots were developed using Western Lightning Plus Chemiluminescence Substrate (Perkin Elmer) and an ImageQuant LAS 4000 (GE). A list of primary antibody is provided below.

| Target | Manufacturer (Product #) | Dilution |
| --- | --- | --- |
| p-AKT | Cell Signaling Technology #4058 | 1:1000 |
| p-ERK1/2 | Cell Signaling Technology #9101 | 1:1000 |
| p-AMPKa | Cell Signaling Technology #2531 | 1:1000 |
| Vinculin | Sigma-Aldrich #V9131 | 1:5000 |
| LC3I/II | Cell Signaling Technology 2775 | 1:1000 |

### RNA Isolation, sequencing and analysis

Tissue samples were homogenized in TRI reagent (Molecular Research Center, Cincinnati, OH, USA) and the aqueous phase was separated and isolated using 1/5 volume of chloroform followed by centrifugation at 12,000 g for 15 minutes. RNA was precipitated with 100% isopropanol and washed with 75% ethanol. RNA pellets were dissolved in HPLC grade water (Sigma Aldrich) and RNA concentrations were determined using a NanoDrop 2000c Spectrophotometer (Thermo Fisher, Waltham, MA, USA). RNA samples were submitted to Novogene for library preparation, by PolyA selection, and sequencing. All samples had a sequencing depth of at least 20 million 150-bp paired-end reads. All differential expressed gene (DEG) analyses were carried out in R (v. 4.0.3) with package DESeq2. Reads were removed from analysis if the expression was less than 0.5 counts/million. Once exclusively DEGs (p-adjusted-value < 0.05) were identified, we used Gene Ontology Enrichment Analysis for identifying predicted functional pathways and upstream regulators.

### cDNA Preparation and Quantitative Real-Time PCR

RNA was reverse transcribed using SuperScript III reverse transcriptase (Invitrogen, 18080044) and the protocol was followed according to the manufacturer’s instructions. cDNA was diluted to 1 µg/µL in water. Each qPCR reaction contained 4 µg cDNA + SYBR Green PCR Master Mix (Invitrogen 4309155) + 12.5 µM primer set. Thermocycler settings were determined using SYBR Green PCR Master Mix protocol. ΔΔCt was calculated using 18S as a normalizer. A list of primers is reported below.

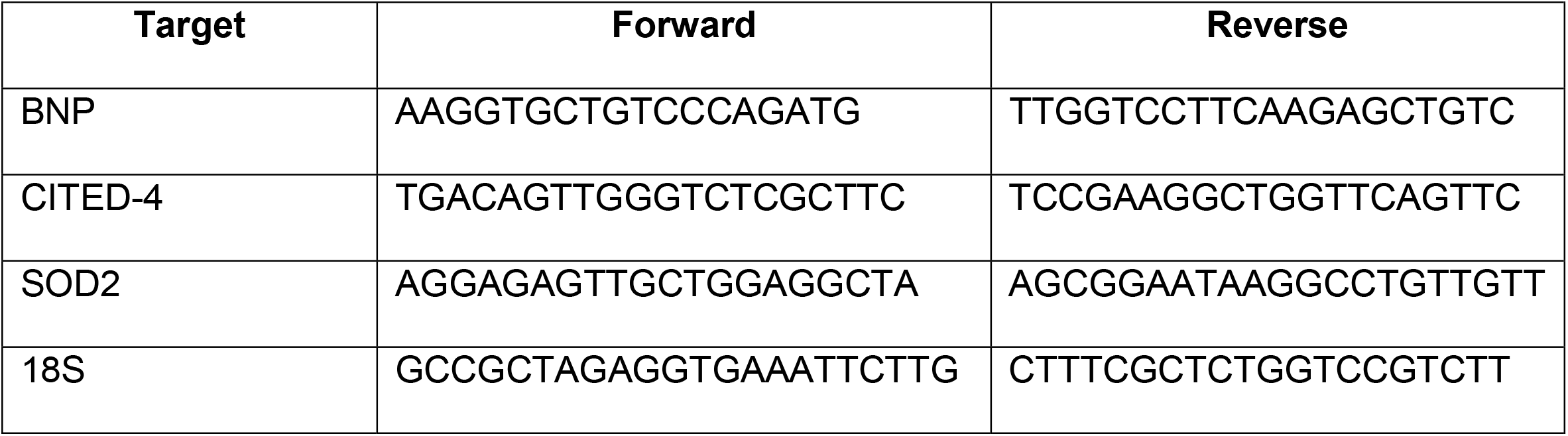

### Statistical analysis

Statistical tests are reported in the figure legend.

## RESULTS

### Age affected cardiac hypertrophy associated with exercise and regression induced by detraining in female mice

In order to understand whether age affects cardiac response to exercise and detraining, female mice were exposed to 28 days of voluntary cage wheel running starting at 5 weeks (young) or 24 weeks (adult) of age and then transferred to a standard housing cage to initiate detraining (Fig 1A). Age-matched sedentary mice were used as controls. Young and adult mice ran similar distances independent of their age (Fig 1B). We defined the following groups: mice at 0 days of detraining (D0), at 3 days of detraining (D3), and at 7 days of detraining (D7). Mice were randomly assigned into groups and no difference in run distances between groups was observed (Fig S1A). Young and adult mice showed significantly different increases in cardiac mass at D0 (for young 22.5 % ± 3.2; for adult 13.7 ± 2.7 [mean ± standard error] (Fig 1C). Adult female mice showed no regression of exercise-induced cardiac hypertrophy after 7 days of detraining, while cardiac mass of young females decreased approximately 10% after 7 days of detraining (Fig 1C). To confirm that the increase of cardiac mass induced by exercise could completely regress with detraining, we measured cardiac mass 28 days after detraining. Both young and adult mice showed complete regression of cardiac mass as compared to sedentary hearts (Fig S1C). We carried out our exercise studies also on young and adult males and confirmed previous work showing sex-dependent differences in exercise performance and cardiac growth in response to exercise^12,13^. Males run less distance on voluntary wheels than females (Figure S1C). As compared to age-matched females, young male mice run on average 21 km less and adult male mice run 127 km less over 4 weeks. Males achieved approximately 5% cardiac hypertrophy after 4 weeks of training, but cardiac mass was not significantly different than sedentary age-matched sedentary male mice (Figure S1C), making it difficult to interpret regression trends and signaling pathway regulation during exercise and regression in male mice. We therefore, focused on female mice from now on.

**Figure 1:**
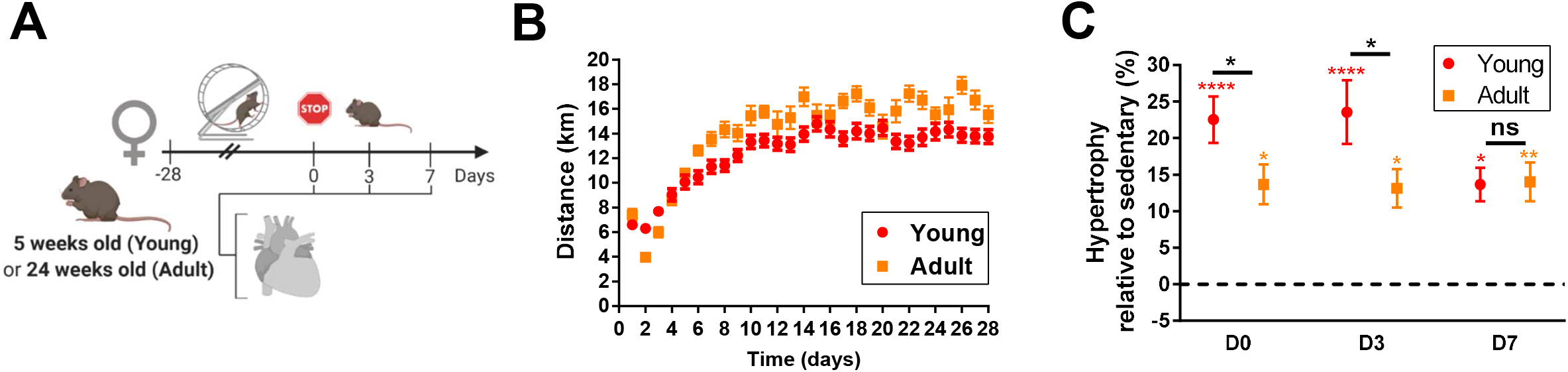
Regression of exercise-induced cardiac hypertrophy. **A)** Scheme of the experimental design. Female mice started running at either 5 or 24 weeks old. Voluntary wheel running was recorded for 28 days. Hearts were collected at 0, 3, and 7 days post running. Sedentary mice were used as controls. **B)** Average running distance (km/day) in young (5-week old) and adult (24-week old) female mice exercised for 28 days in voluntary-wheel cages. **C)** Left ventricle mass normalized by tibia length for each experimental group. Percentages refer to the average relative left ventricle mass for each group compared with the mean cardiac mass of corresponding age-matched sedentary mice. N=14-8 per group. Two-way ANOVA with Tukey's multiple comparisons (\**P<*0.05 **p<0.01, ****p<0.0001). Colored asterisks refer to statistical difference with corresponding age-matched sedentary female mice.

### RNAseq analysis revealed age-dependent response to exercise-induced hypertrophy

Considering the differences in cardiac hypertrophy developed with exercise and regression rate with detraining, we investigated genes expression in mice after 28 days of voluntary wheel running (D0) and 3 days after detraining (D3). We compared gene expression between young and adult female mice. We found that 290 genes were differentially expressed during exercise and 688 gene were differentially expressed 3 days after detraining between young and adult mice (Fig 2A-B). Gene ontology enrichment analysis revealed differences in metabolism and autophagy with exercise and in translation and fatty acid oxidation with detraining (Fig 2C).

**Figure 2:**
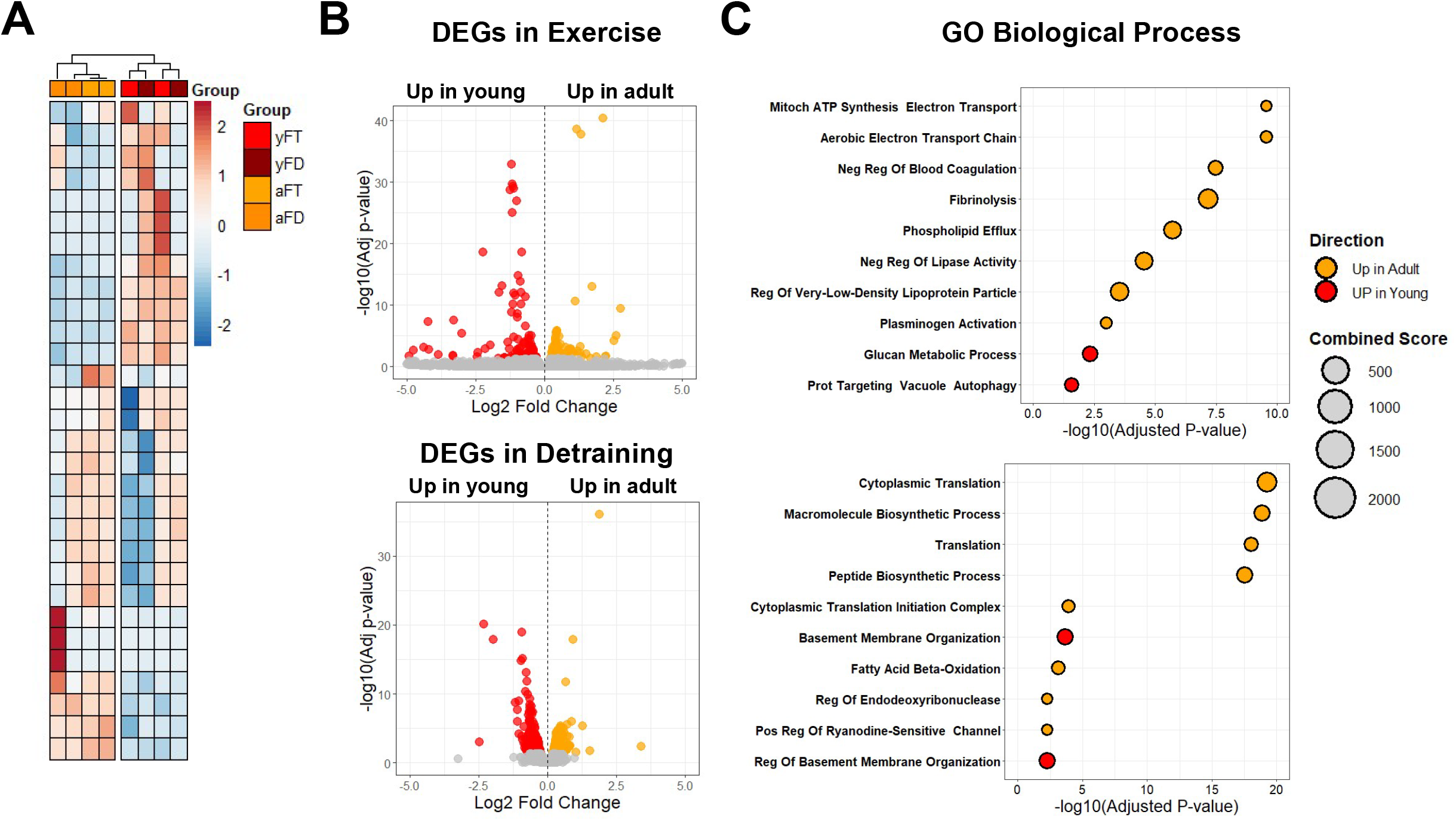
RNAseq analysis of training and detraining in young and adult female mice. **A)** Heatmap of the top 30 differentially-expressed genes in young and adult female mice after 4 weeks of voluntary-wheel running (yFT and aFT, respectively) and 3 days after detraining (yFD and aFD, respectively). **B)** Top, volcano plot of differentially expressed genes (DEGs) comparing young and adult female mice after 4 weeks of running and, below, volcano plot of differentially expressed genes (DEGs) comparing young and adult female mice 3 days after detraining. **C)** Top, gene ontology (GO) enrichment analysis for biological process in young and adult female mice after 4 weeks of voluntary-wheel running and, below, gene ontology (GO) enrichment analysis for biological process in young and adult female mice 3 days after detraining.

### ERK signaling was preferentially activated in young female mice during exercise-induced cardiac hypertrophy

We examined the phosphorylation status of ERK1/2, a key mediator of physiological cardiac hypertrophy. Western blot analysis revealed significantly increased phosphorylation of ERK1/2 in the hearts of young female mice after 28 days of exercise (Figure 3A, red outline), which persisted through day 3 of detraining before returning to baseline levels by day 7. In contrast, adult female hearts showed a significant downregulation in ERK1/2 phosphorylation with exercise (Figure 3A, orange outline). To further investigate the downstream effects of ERK activation, we analyzed the expression of CITED4, a transcriptional regulator implicated in exercise-induced cardiomyocyte proliferation. Consistent with the ERK phosphorylation pattern, CITED4 expression was significantly upregulated in young female hearts following exercise and remained elevated through day 3 of detraining (Figure 3B, red outline). Adult hearts showed only modest, non-significant increases in CITED4 expression (Figure 3B, orange outline). Additionally, we examined the expression of SOD2, an important mitochondrial antioxidant enzyme that helps maintain cardiac function during increased workload. Young female hearts exhibited significant upregulation of SOD2 following exercise, with expression levels gradually decreasing during the detraining period (Figure 3C, red outline). In contrast, adult hearts showed minimal changes in SOD2 expression throughout the study (Figure 3C, orange outline). To exclude that cardiac hypertrophy in adult mice was pathological to some extent, we measured expression of BNP. No changes were found in either group (Figure S2A). Additionally, we tested activation of Akt pathway by measuring phosphorylation of Akt via western blotting. No changes were found in either group (Figure S2B). These findings suggest that preferential activation of the ERK signaling pathway and its downstream targets in young hearts may contribute to the enhanced cardiac hypertrophic response observed in younger animals.

**Figure 3:**
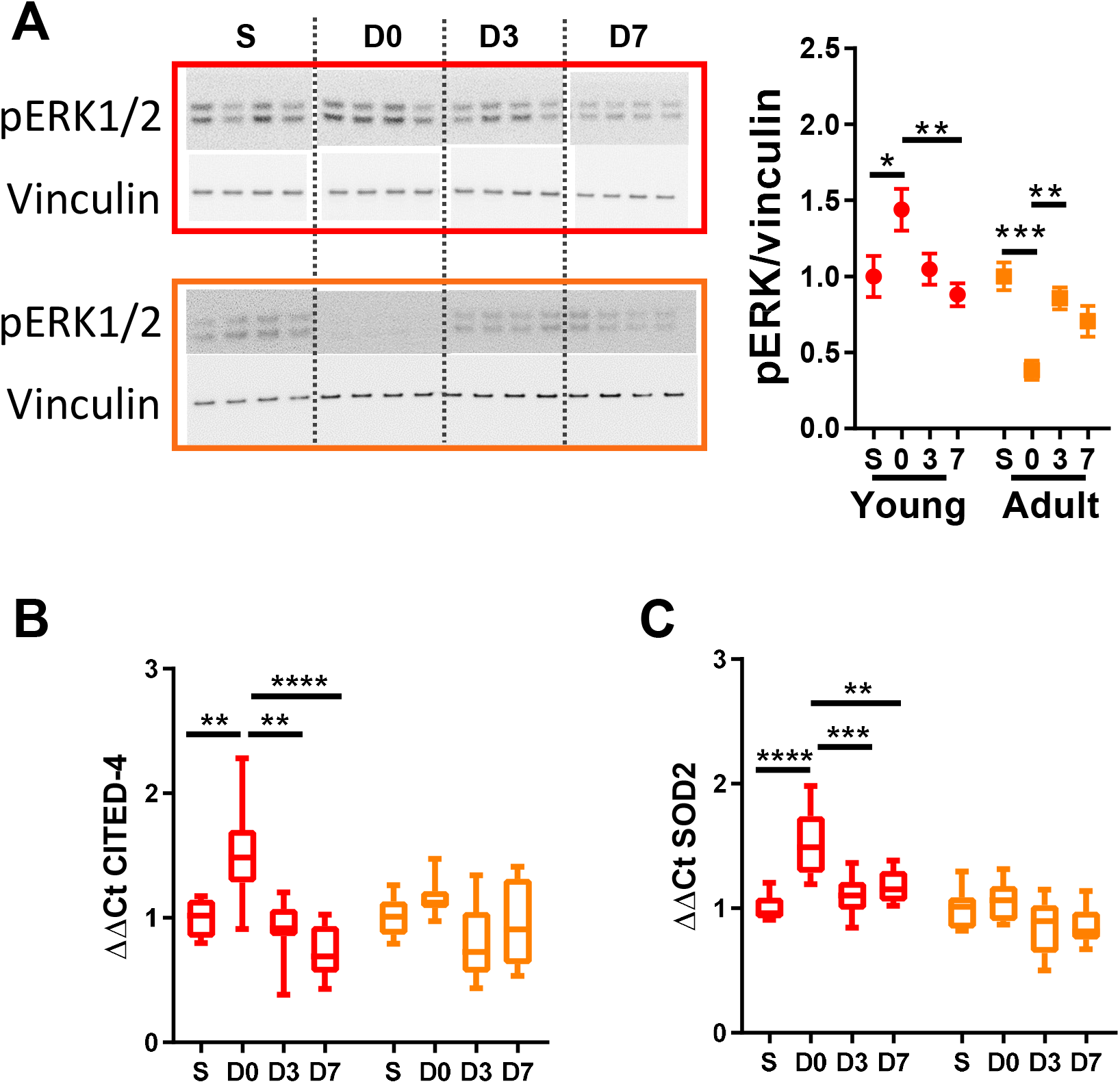
ERK signaling is activated with exercise in young female mice but not adult. A) Western blot of pERK in young female hearts (red outline) and in adult female hearts (orange outline). Quantification on the right. B) Gene expression of CITED4 in young female hearts and in adult female hearts. C) Gene expression of SOD2 in young female hearts and in adult female hearts. N=7-8 per group. One-way ANOVA with Sidak’s multiple comparisons test applied in young and adult group separately (\**P<*0.05, **p<0.01, ***p<0.001, ****p<0.0001).

### Autophagy was preferentially activated during detraining in young female mice

Given the transcriptomic evidence suggesting differential activation of protein degradation pathways, we investigated markers of autophagy. Western blot analysis of LC3, a key autophagy marker, revealed a significant increase in the LC3-II/LC3-I ratio in young female hearts during the detraining period, with peak activation at day 3 of detraining (Figure 4A, red outline). This autophagy activation coincided with the period of rapid cardiac mass regression observed in young mice. In contrast, adult female hearts showed no significant changes in the LC3-II/LC3-I ratio throughout the detraining period (Figure 4A, orange outline). To elucidate the potential upstream regulators of this autophagic response, we examined the phosphorylation status of AMPK, a key sensor of cellular energy status and regulator of autophagy. Young female hearts exhibited significant increases in AMPK phosphorylation during the detraining period, with peak activation at day 3 of detraining (Figure 4B, red outline), mirroring the pattern observed for LC3-II/LC3-I ratio. Adult hearts, however, showed no significant changes in AMPK phosphorylation (Figure 4B, orange outline). Interestingly, comparison of baseline autophagy markers between sedentary young and adult female mice revealed higher basal levels of both the LC3-II/LC3-I ratio and phosphorylated AMPK in young hearts (Figure 4C), suggesting inherently greater autophagy potential in younger animals. These findings indicate that enhanced activation of AMPK-mediated autophagy in young hearts during detraining may contribute to the more rapid regression of exercise-induced cardiac hypertrophy observed in younger animals.

**Figure 4:**
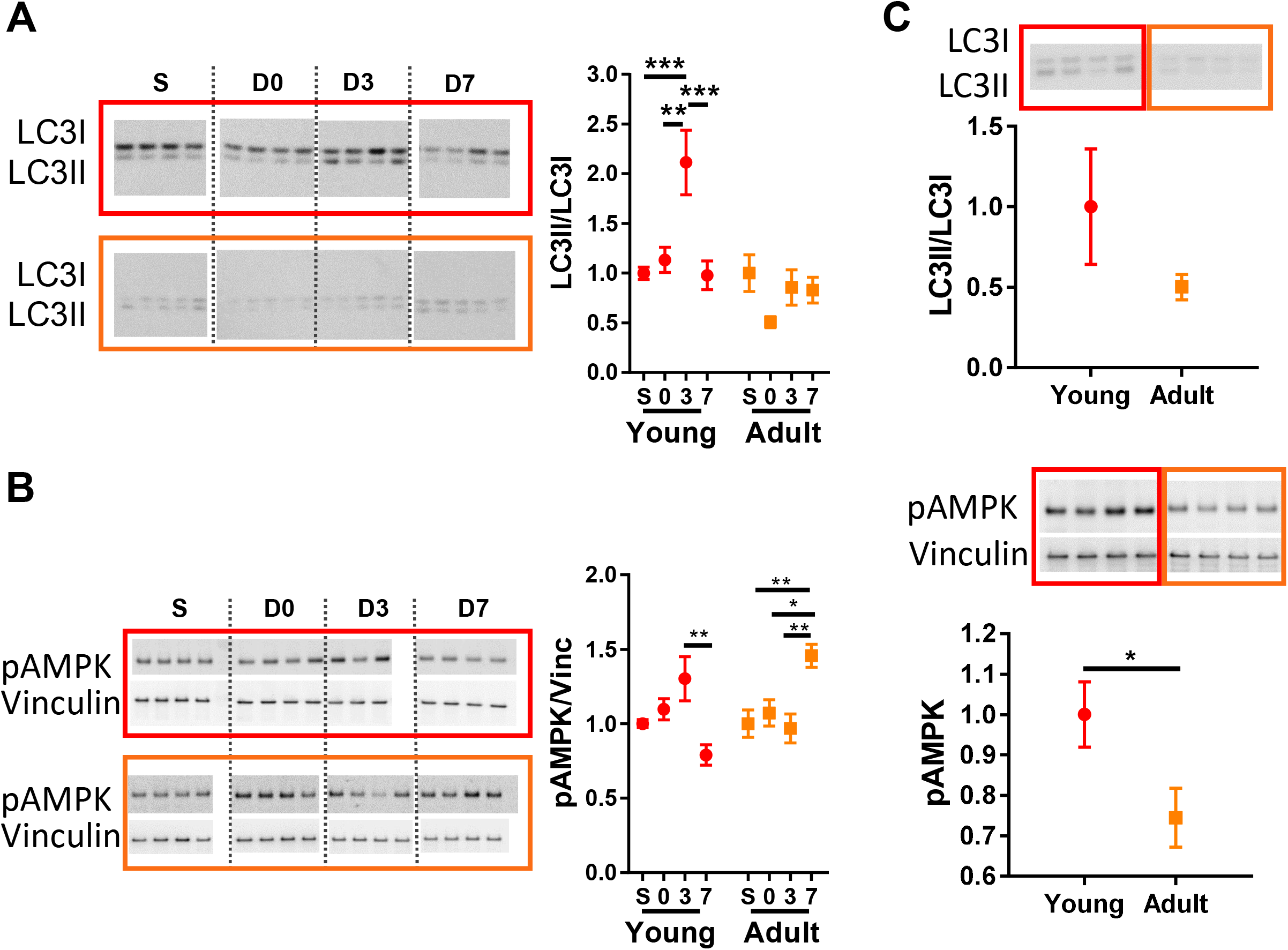
Autophagy is activated in detraining of young female mice but not adult. A) Western blot of LC3I and LC3II in young female mice (red outline) and in adult female mice (orange outline). Quantification on the right. B) Western blot of pAMPK and vinculin in young female mice (red outline) and in adult female mice (orange outline). Quantification on the right. C) Western blot of LC3I and LC3II (above) and pAMPK (below) in sedentary young female mice (red outline) and in adult female mice (orange outline). N=7-8 per group. For A and B, One-way ANOVA with Sidak’s multiple comparisons test (\**P<*0.05, **p<0.01, ***p<0.001). For C, Unpaired t-test (\**P<*0.05).

## DISCUSSION

This study provides valuable insights into the age-dependent differences in cardiac remodeling during exercise and detraining in female mice, highlighting distinct molecular mechanisms that govern cardiac plasticity across the lifespan. Our findings reveal that young female mice exhibit greater cardiac adaptability to both exercise stimulus and its withdrawal, showing enhanced hypertrophic growth and more rapid regression compared to adult counterparts.

The selection of 5-week-old and 24-week-old female mice allowed us to specifically investigate how cardiac adaptability changes from youth to adulthood, rather than focusing on senescence-related cardiac changes. The voluntary wheel running protocol employed in this study offers several advantages over forced exercise paradigms. Unlike treadmill running or swim training, which may induce stress responses that confound physiological adaptations^5^, voluntary wheel running more accurately reflects natural rodent behavior and eliminates stress-related variables. The 28-day exercise period followed by variable detraining intervals (3, 7, and 28 days) provided a comprehensive timeline to observe both the development and regression of cardiac hypertrophy. Importantly, both age groups demonstrated similar running distances (Figure 1B and S1A), eliminating exercise volume as a confounding variable and allowing direct comparison of age-dependent molecular responses to equivalent exercise stimuli.

Our focus on female mice was motivated by the established sexual dimorphism in exercise-induced cardiac remodeling^12–14,30^ and our results on male mice. Previous studies have demonstrated that female mice typically exhibit more pronounced cardiac hypertrophy in response to exercise compared to males, with increases of 15-25% versus 5-9% in cardiac mass, respectively^12^. Our findings of 23% hypertrophy in young females and 15% in adult females (Figure 1C and S1B) align with these sex-specific patterns while revealing additional age-dependent nuances. We also confirmed reduced exercise performance and cardiac hypertrophy in both young and adult male mice (Figure S1D-F).

It is worth noting that the natural tendency of mice toward physical activity suggests that the sedentary condition may represent a departure from their physiological norm. This raises the possibility that the sedentary state could be considered the “pathological” condition, rather than the exercise state representing a deviation from baseline. This perspective aligns with previous considerations in the general animal research literature^31,32^ and in the cardiac research literature^4,9,33^. Evolutionary biology concepts also suggest that regular physical activity represents the default physiological state for mammals^34^.

The marked difference in hypertrophic response between young and adult mice (23% versus 15% increase in left ventricular mass) despite similar running distances indicates inherent age-related differences in cardiac plasticity. Even more striking was the differential regression pattern, with young mice demonstrating significant cardiac mass reduction by day 7 of detraining, while adult mice maintained their exercise-induced hypertrophy through this time point (Figure 1C). The complete regression observed in both age groups by day 28 confirms the physiological nature of the hypertrophic response (Figure S1C).

This age-dependent regression pattern has important clinical implications. In human athletes, detraining serves as a diagnostic tool to differentiate physiological “athlete's heart” from pathological cardiomyopathies^35^. Regression of athletic left ventricular hypertrophy typically occurs within 1-3 months of detraining in humans and is characterized by decreases in the intracellular myocardial compartment without changes in the extracellular matrix^36^. The delayed regression observed in our adult mice raises questions about whether the slower adaptive response represents reduced cardiac flexibility or perhaps a more efficient retention of beneficial adaptations. The eventual complete regression in both age groups by day 28 suggests that while kinetics differ, the fundamental physiological nature of exercise-induced hypertrophy is preserved across these age groups.

The transcriptomic analysis revealed substantial age-dependent differences in gene expression profiles during both exercise (290 differentially expressed genes) and early detraining (688 differentially expressed genes) (Figure 2A-B). These findings underscore the molecular diversity underlying apparently similar phenotypic changes. The enrichment of metabolic and autophagy pathways during exercise and translation and fatty acid oxidation during detraining (Figure 2C) suggests fundamentally different adaptation mechanisms between young and adult hearts. These distinct molecular signatures may explain the differential magnitude and kinetics of cardiac remodeling observed between age groups. The greater number of differentially expressed genes during early detraining (D3) compared to the exercise period (D0) is particularly intriguing, suggesting that cardiac regression may involve more complex and active molecular reprogramming than previously appreciated, rather than simply representing the cessation of hypertrophic signaling. This supports the concept that cardiac regression following exercise is an active process requiring coordinated molecular events^28^, particularly in younger animals where regression occurs more rapidly.

In our work, ERK1/2 signaling is robustly activated in young female hearts following exercise and persisted through early detraining (Fig 3A). Adult mice showed a significant downregulation of ERK1/2 phosphorylation. This differential activation pattern was mirrored in downstream targets CITED4 and SOD2 (Figure 3B-C), both established mediators of physiological cardiac adaptation.

CITED4 has been implicated in exercise-induced cardiomyocyte proliferation^9^, suggesting that enhanced CITED4 expression in young hearts may contribute to their greater hypertrophic capacity. Similarly, the robust upregulation of SOD2 in young but not adult hearts indicates enhanced mitochondrial antioxidant defense, which is crucial for maintaining cardiac function during increased workload^37^. These findings collectively suggest that young hearts possess greater capacity to activate adaptive signaling pathways in response to exercise stimuli.

Adult female mice specifically inhibit ERK signaling but still achieved significant hypertrophy (13.7% increase in mass). This is in line with previous work showing that ERK is not essential for cardiac hypertrophy in response to exercise^38^. It appears that young female mice activate ERK pathway to enhance their cardiac hypertrophy with exercise. Other hypertrophic pathways are likely involved in both young and adult mice. Our results excluded activation of pathological hypertrophy (Figure S2A) and Akt signaling (Figure S2B). Nonetheless, this age-dependent signaling switch merits further investigation, as it may reveal novel therapeutic targets for enhancing cardiac adaptability across the lifespan.

The differential activation of autophagy between young and adult hearts during detraining (Figure 4A) provides a compelling mechanism for the observed differences in regression kinetics. The significant increase in LC3-II/LC3-I ratio and AMPK phosphorylation (Figure 4B) exclusively in young hearts during early detraining (D3) coincided with the period of rapid cardiac mass regression. This temporal alignment suggests that enhanced autophagy-mediated protein degradation contributes to the accelerated regression observed in younger animals.

The higher baseline levels of autophagy markers in sedentary young versus adult hearts (Figure 4C) further indicates an age-dependent reduction in autophagy potential. This diminished capacity for protein quality control and cellular remodeling in adult hearts may explain their slower regression following exercise cessation. Given that efficient autophagy is essential for cellular homeostasis and adaptation to stress^39^, this finding has broader implications for understanding age-related declines in cardiac adaptability.

The AMPK-mediated activation of autophagy in young hearts during detraining represents a valuable finding in the context of physiological cardiac regression. AMPK serves as a cellular energy sensor that responds to changes in the AMP/ATP ratio, suggesting that early detraining may create a transient energy imbalance that triggers adaptive autophagy. This process likely facilitates the efficient removal of surplus cellular components acquired during hypertrophy, allowing rapid return to baseline cardiac dimensions. It would be valuable to compare mechanisms of regression between physiological and pathological cardiac hypertrophy^40^.

Our findings have important implications for understanding exercise-induced cardiac preconditioning. Previous research has demonstrated that exercise-induced hypertrophy (swimming) increases resistance to pathological stress through what has been termed “anti-hypertrophic memory"^27^. This protective effect is mediated through a signaling pathway involving Mhrt779/Brg1/Hdac2/p-Akt/p-GSK3β, which attenuates pathological hypertrophic responses to subsequent stressors^27^. The age-dependent differences in molecular adaptation observed in our study suggest that this preconditioning effect may vary with age. The robust ERK activation and downstream signaling in young hearts may contribute to more effective preconditioning, potentially conferring greater protection against future pathological stressors. Conversely, the diminished signaling response in adult hearts may indicate reduced preconditioning efficacy. However, the more persistent hypertrophy during detraining in adult hearts could alternatively reflect a more stable form of cardiac adaptation, potentially providing sustained protection despite less dynamic molecular responses.

The concept of “anti-hypertrophic memory” established by previous studies may operate differently across the lifespan. Young hearts, with their enhanced signaling responses and autophagy capacity, may develop more flexible but transient protective mechanisms. Adult hearts, with their more persistent but attenuated molecular responses, may establish more stable but less adaptable protective mechanisms. Understanding these age-dependent differences in cardiac preconditioning could inform more nuanced exercise recommendations across the lifespan.

Several limitations of the current study warrant consideration. Future studies should include multiple age points provide a more comprehensive understanding of cardiac adaptation throughout the lifespan. Moreover, our investigation focused primarily on ERK signaling and autophagy pathways, leaving other potential mechanisms unexplored. The transcriptomic analysis revealed numerous differentially expressed genes that merit further investigation, particularly those involved in metabolism and protein translation. Assessment of other canonical hypertrophic pathways would provide a more complete picture of age-dependent cardiac adaptation. Finally, functional assessment of cardiac performance through echocardiography or pressure-volume analysis would complement our structural and molecular findings, revealing whether the observed age-dependent differences in cardiac remodeling translate to functional advantages or disadvantages.

This study demonstrates that cardiac adaptation to exercise and detraining follows distinct molecular trajectories in young versus adult female mice. Young hearts exhibit enhanced plasticity characterized by robust ERK signaling during hypertrophy and elevated autophagy during regression, resulting in more pronounced hypertrophic growth and more rapid regression. These age-dependent differences in cardiac plasticity have important implications for understanding the cardiovascular benefits of exercise across the lifespan and developing age-appropriate exercise recommendations.

## ACKNOWLEDGMENT

We thank Amanda Wacker and Deanna Muehleman for their assistance. This research was supported by the Joyce and Dick Brown endowed chair at University of Colorado Boulder and by the DZHK (German Centre for Cardiovascular Research), partner site Berlin (Project: 81X3100222).

