## Supplemental Figure 1 for "Age-Dependent Mechanisms of Cardiac Hypertrophy Regression Following Exercise in Female Mice"

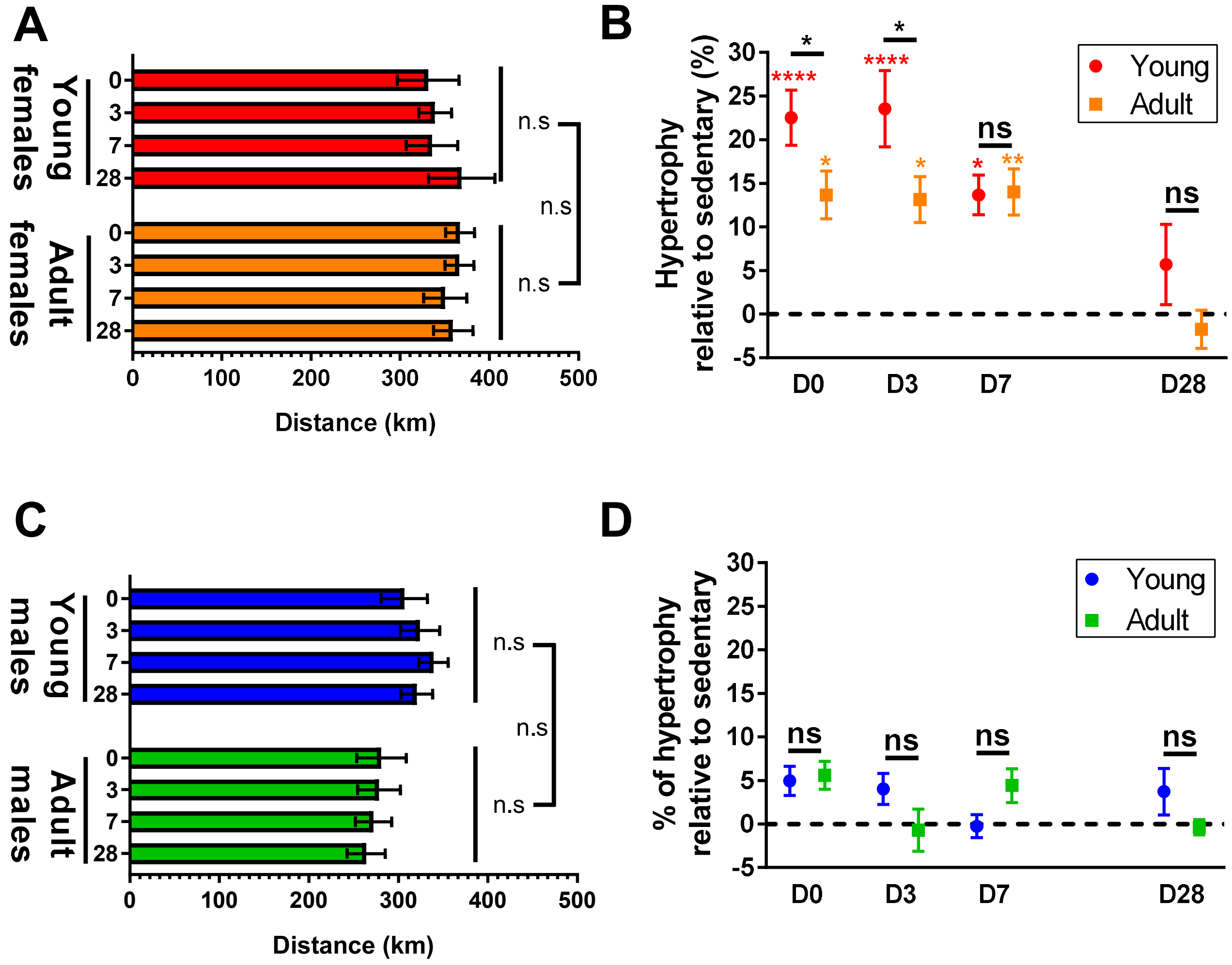


**Figure S1:** A) total distance run by each group. Female animals were randomly assigned to the groups. N=7-8. One way ANOVA with Tukey's multiple comparisons test applied to compare every group. B) Left ventricle mass normalized by tibia length for each experimental group. Percentages refer to the average relative left ventricle mass for each group compared with the mean cardiac mass of corresponding age-matched sedentary female mice. N=4-14 per group. Two-way ANOVA with Tukey's multiple comparisons (*$P$<0.05 **p<0.01, ****p<0.0001). Colored asterisks refer to statistical difference with corresponding age-matched sedentary female mice. C) total distance run by each group. Male animals were randomly assigned to the groups. N=6-10. One way ANOVA with Tukey's multiple comparisons test applied to compare every group. D) Left ventricle mass normalized by tibia length for each experimental group. Percentages refer to the average relative left ventricle mass for each group compared with the mean cardiac mass of corresponding age-matched sedentary male mice. N=7-9 per group. Two-way ANOVA with Tukey's multiple comparisons test.
