## Supplemental Figure 2 for "Age-Dependent Mechanisms of Cardiac Hypertrophy Regression Following Exercise in Female Mice"

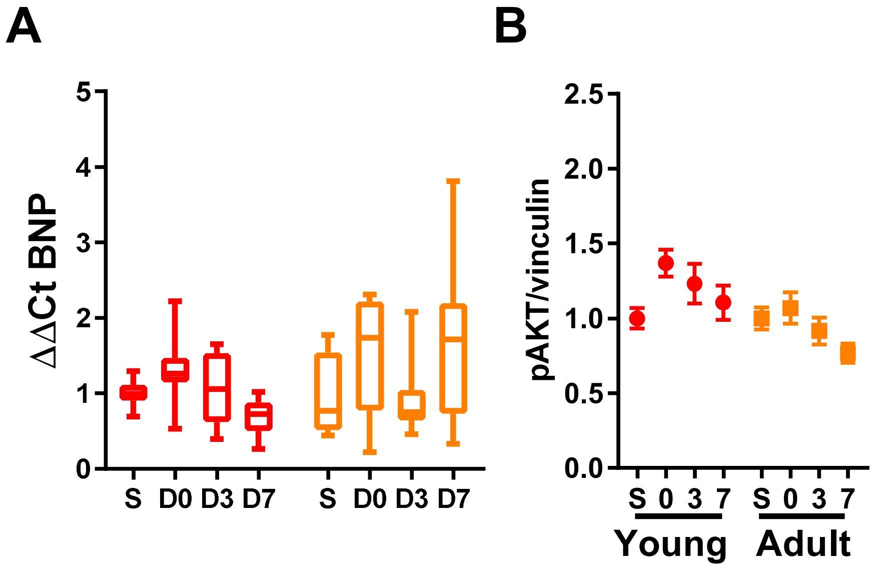


**Figure S2:** A) Gene expression of BNP in young female hearts and in adult female hearts. N=7-8 per group. Kruskal-Wallis test with Dunn's multiple comparisons test applied in young and adult group separately. B) Western blot pAKT normalized by vinculin in young and adult female mice. N=7-8 per group. One-way ANOVA with Sidak's multiple comparisons test applied in young and adult group separately.
